## Supplemental Information for "Beyond the Active Site: The addition of a remote loop reveals a new complex biological function for chitinase enzymes"

**Supplementary Table 1 Primers used in this study**

| **Primers** | **Sequence 5' to 3'** |
|---|---|
| T7 | TAATACGACTCACTATAGGG |
| T7_terminator | GCTAGTTATTGCTCAGCGG |
| Linear_vector_F | TAGTGAGTCGTATTAATTTCG |
| Linear_vector_R | TGAGCAATAACTAGCATAACCCCTTG |
| **Anc4** | |
| +Loop I_F | GCCCTTCGCAGGGCACGCGCCACTATTACGGTTCGGGAACAG |
| +Loop I_R | GGCGCGTGCCCTGCGAAGGGCTTTTACACGTATGACGCG |
| +Loop II_F | CCACGCGTAAGGGCCATCCGGTGCGGTCGGCCATCCACCACTAGTTTCGTGACTCAC |
| +Loop II_R | GGTGGATGGCCGACCGCACCGGATGGCCCTTACGCGTGGGGCTTGGTATATATTGAGGAAATTAATC |
| E57Q_F | ACTAGTTTGGTGACTCACATGTGC |
| E57Q_R | AGTCACCAAACTAGTGGCTTGG |
| **Anc4+LoopII** | |
| E57Q_R | AGTCACCAAACTAGTGGTGGATGGC |
| p12K/n13H_F | ATTACGGTGTTTGAACAGTTGGTCGAACAT |
| p12K/n13H_R | CAACTGTTCAAACACCGTAATAGTTTTTAC |
| s58T_F | TCCACCAGTAGTTTCGTGACTCACATG |
| s58T_R | GAAACTACTGGTGGATGGCCGACCGCAC |
| n193G/y194F/d197R_F | GAATCTTTTATAAAATCCCACACGATTCTC |
| n193G/y194F/d197R_R | GTGGGATTTTATAAAAGATTCTGCAATCAG |
| **Anc5** | |
| ΔLoop I_F | TGTAGAAGTCGTTGCGGTGTTTCAA |
| ΔLoop I_R | CAACGACTTCTACACATACGATGCG |
| ΔLoop II_F | AGTAGCCTGTAGTCTCATGACTCGTC |
| ΔLoop I_R | GACTACAGGCTACTGTTTTCTGAAAGAG |
| E64Q_F | TGTAGTCTGATGACTCGTCTGACC |
| E64Q_R | AGTCATCAGACTACAGGTGGATGGC |
| P204n/D205p_F | TTGTGCCGGGTTTCCTTTCCCGCACTC |
| P204n/D205p_R | GGGAAAGGAAACCCGGCACAAGTAGAGGAC |
| **Anc5ΔLoopII** | |
| E64Q_R | AGTCATCAGACTACAGGCTACTGTTTTC |

**Supplementary Table 2 Experimental Details for crystallization of Anc4 and Anc5**

| **Structure** | Anc4 | Anc5 |
|---|---|---|
| **PDB code** | 8HNE | 8HNF |
| **Buffer** | 5 mM Tris-HCl pH 8.0 | 5 mM Tris-HCl pH 8.0 |
| **Crystallization** | Hanging drop, 1 μL of Anc4 (10 mg/mL) + 1 μL of 15% (w/v) PEG mono ether 2,000, 0.1 M MES pH 6.0 | Hanging drop, 1 μL of Anc5 (5.67 mg/mL) + 1 μL of 27% (w/v) PEG mono ether 2,000, 0.1 M MES pH 6.0 |
| **Cryoprotectant** | 40% (w/v) PEG monomethyl ether 2,000, 0.1 M MES pH 6.0 | 40% (w/v) PEG monomethyl ether 2,000, 0.1 M MES pH 6.0 |
| **Wavelength (Å)** | 1.000 | 1.000 |
| **Beamline** | SPring-8 BL45XU | SPring-8 BL41XU |
| **Data processing** | KAMO, XDS, XSCALE | XDS, Aimless |
| **MR model** | AlphaFold2 | AlphaFold2 |
| **Refinement** | The model building was performed with *Coot*. The model structure was refined by *Coot* and Phaser. | The model building was performed with *Coot*. The model structure was refined by *Coot* and Phaser. |

**Supplementary Table 3 Experimental Details for crystallization of Anc4+Loop II mutants**

| | | |
|---|---|---|
| **Structure**<br>**PDB code** | Anc4+Loop II+p12K/n13H | Anc4+Loop II+p12K/n13H/s58T/n193G/y194F/d197R |
| **Buffer** | 10 mM Sodium acetate,<br>150 mM NaCl pH 5.0 | 10 mM Sodium acetate,<br>150 mM NaCl pH 5.0 |
| **Crystallization** | Sitting drop, 1 μL of Anc4+Loop II+p12K/n13H (6.00 mg/mL) + 1 μL of 25% (w/v) polyethylene glycol 3,350, 0.1 M Bis-Tris, pH 6.5 | Sitting drop, 1 μL of Anc4+Loop II+p12K/n13H/s58T/n193G/y194F/d197R (3.80 mg/mL) + 1 μL of 25% (w/v) polyethylene glycol 3,350, 0.1 M Bis-Tris, pH 6.5 |
| **Cryoprotectant** | 30% (w/v) polyethylene glycol 3,350, 0.1 M Bis-Tris, pH 6.5 | 30% (w/v) polyethylene glycol 3,350, 0.1 M Bis-Tris, pH 6.5 |
| **Wavelength (Å)** | 1.000 | 1.000 |
| **Beamline** | SPring-8 BL32XU | SPring-8 BL32XU |
| **Data processing** | KAMO, XDS, XSCALE | KAMO, XDS, XSCALE |
| **MR model** | AlphaFold2 | AlphaFold2 |
| **Refinement** | The model building was performed with *Coot*. The model structure was refined by *Coot* and Phaser. | The model building was performed with *Coot*. The model structure was refined by *Coot* and Phaser. |

**Supplementary Table 4 Data collection and refinement statistics for Anc4 and Anc5**

| Structure | Anc4 | Anc5 |
|---|---|---|
| PDB code | 8HNE | 8HNF |
| **Data collection** | | |
| Space group | $P2_12_12_1$ | $P12_11$ |
| Cell dimensions | | |
| *a*, *b*, *c* (Å) | 32.29, 76.15, 79.18 | 36.89, 73.43, 39.28 |
| α, β, γ (°) | 90.00, 90.00, 90.00 | 90.00, 99.965, 90.00 |
| Resolution (Å) | 39.47 – 1.13 (1.17-1.13)* | 38.69 – 1.571 (1.627-1.571) |
| $R_{merge}$ | 0.2433 (0.325) | 0.0658 (0.484) |
| $I / \sigma I$ | 4.14 (2.54) | 10.36 (1.18) |
| Completeness (%) | 99.62 (96.37) | 99.54 (96.92) |
| Redundancy | 2.0 (2.0) | 2.0 (2.0) |
| **Refinement** | | |
| Resolution (Å) | 39.47 – 1.13 | 38.69 – 1.57 |
| No. reflections | 73261 (6976) | 28613 (2773) |
| $R_{work}$ / $R_{free}$ (%) | 24.07 / 25.05 | 17.60 / 20.21 |
| No. atoms | | |
| Protein | 1700 | 1881 |
| Ligand/ion | 91 | 66 |
| Water | 214 | 86 |
| *B*-factors | | |
| Protein | 8.82 | 25.15 |
| Ligand/ion | 14.53 | 38.25 |
| Water | 17.40 | 30.73 |
| R.m.s. deviations | | |
| Bond lengths (Å) | 0.007 | 0.010 |
| Bond angles (°) | 1.09 | 1.08 |

*Values in parentheses are for highest-resolution shell.

**Supplementary Table 5 Data collection and refinement statistics for Anc4+Loop II mutants**

| Structure<br>PDB code | Anc4+LoopII+p12K/n13H<br>(8X2V) | Anc4+LoopII+p12K/n13H/s58T/ n193G/y194F/d197R<br>(8X2W) |
|---|---|---|
| **Data collection** | | |
| Space group | $P22_12_1$ | $P22_12_1$ |
| Cell dimensions | | |
| *a*, *b*, *c* (Å) | 37.29, 69.83, 70.27 | 37.32, 69.59, 70.22 |
| α, β, γ (°) | 90.00, 90.00, 90.00 | 90.00, 90.00, 90.00 |
| Resolution (Å) | 49.53 – 1.29 (1.336-1.29)* | 49.43 – 1.40 (1.45-1.40) |
| $R_{merge}$ | 18.6 (162.2) | 234.1 (-99.9) |
| $I$ / σ$I$ | 12.87 (2.13) | 7.00 (0.00) |
| Completeness (%) | 100.0 (100.0) | 99.99 (100.0) |
| Redundancy | 33.1 (26.4) | 55.3 (54.2) |
| **Refinement** | | |
| Resolution (Å) | 39.47 – 1.13 | 38.69 – 1.57 |
| No. reflections | 47013 (4640) | 36817 (3631) |
| $R_{work}$ / $R_{free}$ (%) | 17.1 / 18.6 | 18.64 / 20.41 |
| No. atoms | | |
| Protein | 1818 | 1799 |
| Ligand/ion | 0 | 0 |
| Water | 134 | 118 |
| *B*-factors | | |
| Protein | 14.60 | 23.24 |
| Ligand/ion | 0 | 0 |
| Water | 18.95 | 28.35 |
| R.m.s. deviations | | |
| Bond lengths (Å) | 0.007 | 0.009 |
| Bond angles (°) | 0.96 | 1.06 |

*Values in parentheses are for highest-resolution shell.

**Supplementary Table 6 Effects of mutations that destabilize a loop region on enzymatic activity.**

| variants | Hydrolytic activity (U*/mol) ×$10^9$ | Antifungal activity ($IC_{50}$, μM) |
|---|---|---|
| Anc5 | 0.85 | 16.9 ± 1.4 |
| Anc5ΔLoopI | 1.28 | 17.2 ± 1.9 |
| Anc5 P204n/D205p | 0.92 | 15.4 ± 0.2 |
| Anc5ΔLoopI P204n/D205p | 1.32 | 19.4 ± 2.7 |

One unit of activity is defined as the enzyme activity that produced one μmol of GlcNAc per minute at 37°C.

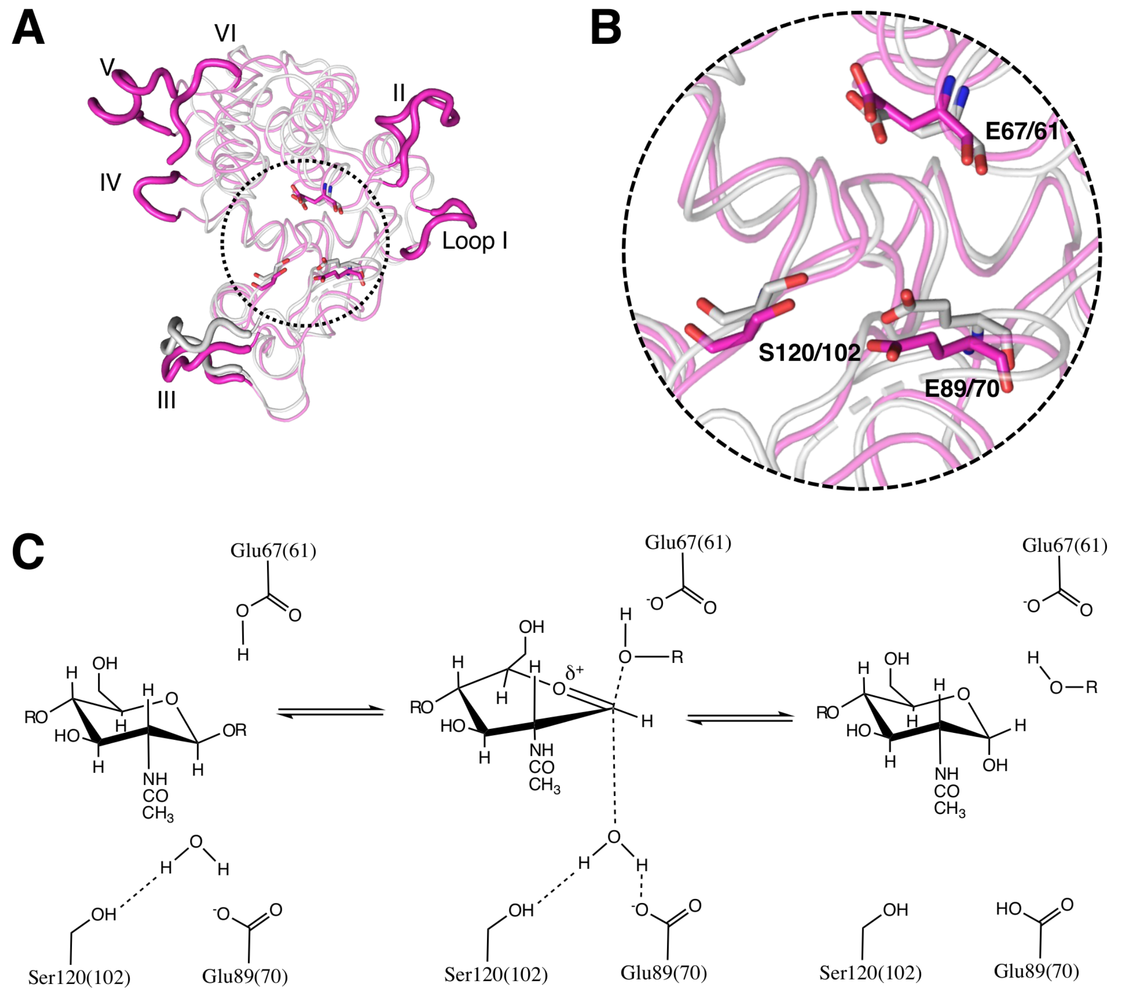

**Figure S1. Conserved catalytic resides and the catalytic mechanism of GH19 chitinase.**

A, Superimposition of loopless (PDB ID:3WH1 with modification of replacing Ala 61 with Glu) and loopful (PDB ID:4J0L with modification of replacing Gln 67 with Glu) GH19 chitinases are shown respectively in gray and magenta. The dashed circle indicates the catalytic center. Highlighted in stick the catalytic residues. B, The enlarged view of the catalytic center shown in A. Two glutamate residues and one serine residue are shown in stick. C, The catalytic mechanism of hydrolysis of beta-1,4-glycosidic linkage proposed by Brameld and Goddard. The numbers of the two glutamate residues and one serine residue are according to loopful GH19; those in parentheses are according to loopless GH19.

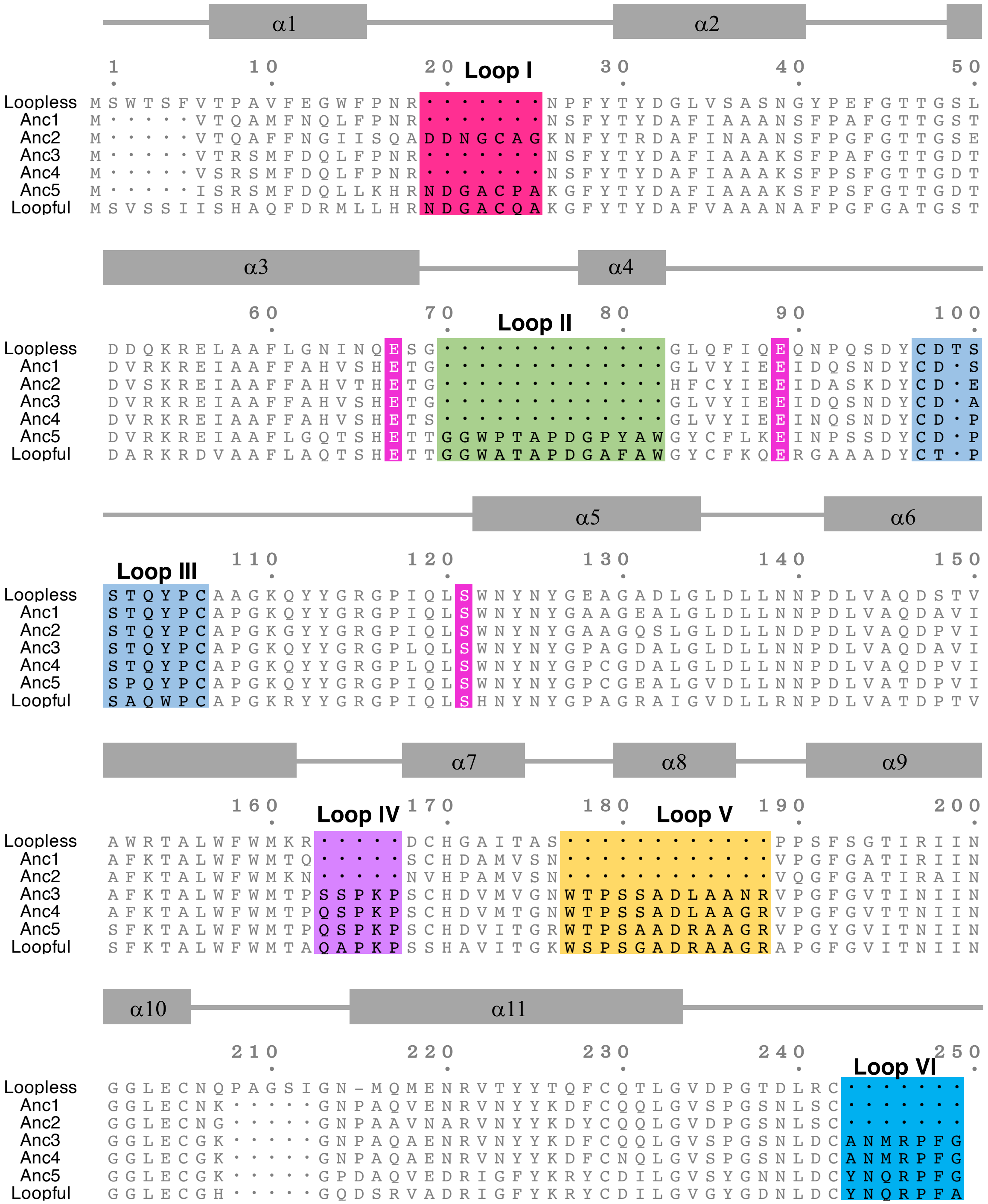

α1
α2
1
10
20
Loop I
30
40
50
Loopless MSWTSFVTPAVFEGWFPNR·······NPFYTYDGLVSASNGYPEFGTTGSL
Anc1 M·····VTQAMFNQLFPNR·······NSFYTYDAFIAAANSFPAFGTTGST
Anc2 M·····VTQAFFNGIISQADDNGCAGKNFYTRDAFINAANSFPGFGTTGSE
Anc3 M·····VTRSMFDQLFPNR·······NSFYTYDAFIAAAKSFPAFGTTGDT
Anc4 M·····VSRSMFDQLFPNR·······NSFYTYDAFIAAAKSFPSFGTTGDT
Anc5 M·····ISRSMFDQLLKHRNDGACPAKGFYTYDAFIAAAKSFPSFGTTGDT
Loopful MSVSSIISHAQFDRMLLHRNDGACQAKGFYTYDAFVAAANAFPGFGATGST
α3
α4
60
70
Loop II
80
90
100
Loopless DDQKRELAAFLGNINQESG·············GLQFIQEQNPQSDYCDTS
Anc1 DVRKREIAAFFAHVSHETG·············GLVYIEEIDQSNDYCD·S
Anc2 DVSKREIAAFFAHVTHETG·············HFCYIEEIDASKDYCD·E
Anc3 DVRKREIAAFFAHVSHETG·············GLVYIEEIDQSNDYCD·A
Anc4 DVRKREIAAFFAHVSHETS·············GLVYIEEINQSNDYCD·P
Anc5 DVRKREIAAFLGQTSHETTGGWPTAPDGPYAWGYCFLKEINPSSDYCD·P
Loopful DARKRDVAAFLAQTSHETTGGWATAPDGAFAWGYCFKQERGAAADYCT·P
α5
α6
Loop III
110
120
130
140
150
Loopless STQYPCAAGKQYYGRGPIQLSWNYNYGEAGADLGLDLLNNPDLVAQDSTV
Anc1 STQYPCAPGKQYYGRGPIQLSWNYNYGAAGEALGLDLLNNPDLVAQDAVI
Anc2 STQYPCAPGKGYYGRGPIQLSWNYNYGAAGQSLGLDLLNDPDLVAQDPVI
Anc3 STQYPCAPGKQYYGRGPLQLSWNYNYGPAGDALGLDLLNNPDLVAQDAVI
Anc4 STQYPCAPGKQYYGRGPLQLSWNYNYGPCGDALGLDLLNNPDLVAQDPVI
Anc5 SPQYPCAPGKQYYGRGPIQLSWNYNYGPCGEALGVDLLNNPDLVATDPVI
Loopful SAQWPCAPGKRYYGRGPIQLSHNYNYGPAGRAIGVDLLRNPDLVATDPTV
α7
α8
α9
160
Loop IV
170
180
Loop V
190
200
Loopless AWRTALWFWMKR·····DCHGAITAS············PPSFSGTIRIIN
Anc1 AFKTALWFWMTQ·····SCHDAMVSN············VPGFGATIRIIN
Anc2 AFKTALWFWMKN·····NVHPAMVSN············VQGFGATIRAIN
Anc3 AFKTALWFWMTPSSPKPSCHDVMVGNWTPSSADLAANRVPGFGVTINIIN
Anc4 AFKTALWFWMTPQSPKPSCHDVMTGNWTPSSADLAAGRVPGFGVTTNIIN
Anc5 SFKTALWFWMTPQSPKPSCHDVITGRWTPSAADRAAGRVPGYGVITNIIN
Loopful SFKTALWFWMTAQAPKPSSHAVITGKWSPSGADRAAGRAPGFGVITNIIN
α10
α11
210
220
230
240
250
Loop VI
Loopless GGLECNQPAGSIGN-MQMENRVTYYTQFCQTLGVDPGTDLRC·······
Anc1 GGLECNK·····GNPAQVENRVNYYKDFCQQLGVSPGSNLSC·······
Anc2 GGLECNG·····GNPAAVNARVNYYKDYCQQLGVDPGSNLSC·······
Anc3 GGLECGK·····GNPAQAENRVNYYKDFCQQLGVSPGSNLDCANMRPFG
Anc4 GGLECGK·····GNPAQAENRVNYYKDFCNQLGVSPGSNLDCANMRPFG
Anc5 GGLECGK·····GPDAQVEDRIGFYKRYCDILGVSYGNNLDCYNQRPFG
Loopful GGLECGH·····GQDSRVADRIGFYKRYCDILGVGYGDNLDCYNQRPFA

**Figure S2. A multiple sequence alignment of two extant GH19 chitinases and five inferred ancestral proteins.**

Multiple sequence alignment was generated by MAFFT and Aliview. Residue numbering and the schematic representation of secondary structures are based on loopful type GH19 chitinase from *Secale cereale* (Uniprot: Q9FRV0; residues 24-266, PDB ID: 4J0L). Loopless indicates the sequence of GH19 chitinase from *Gemmabryum coronatum* (Uniprot: A9ZSX9; residues 25-228). They differ in loop regions, named I to VI highlighted red, green, light blue, purple, orange, and cyan, respectively. Two glutamate residues and one serine residue are shown in magenta.

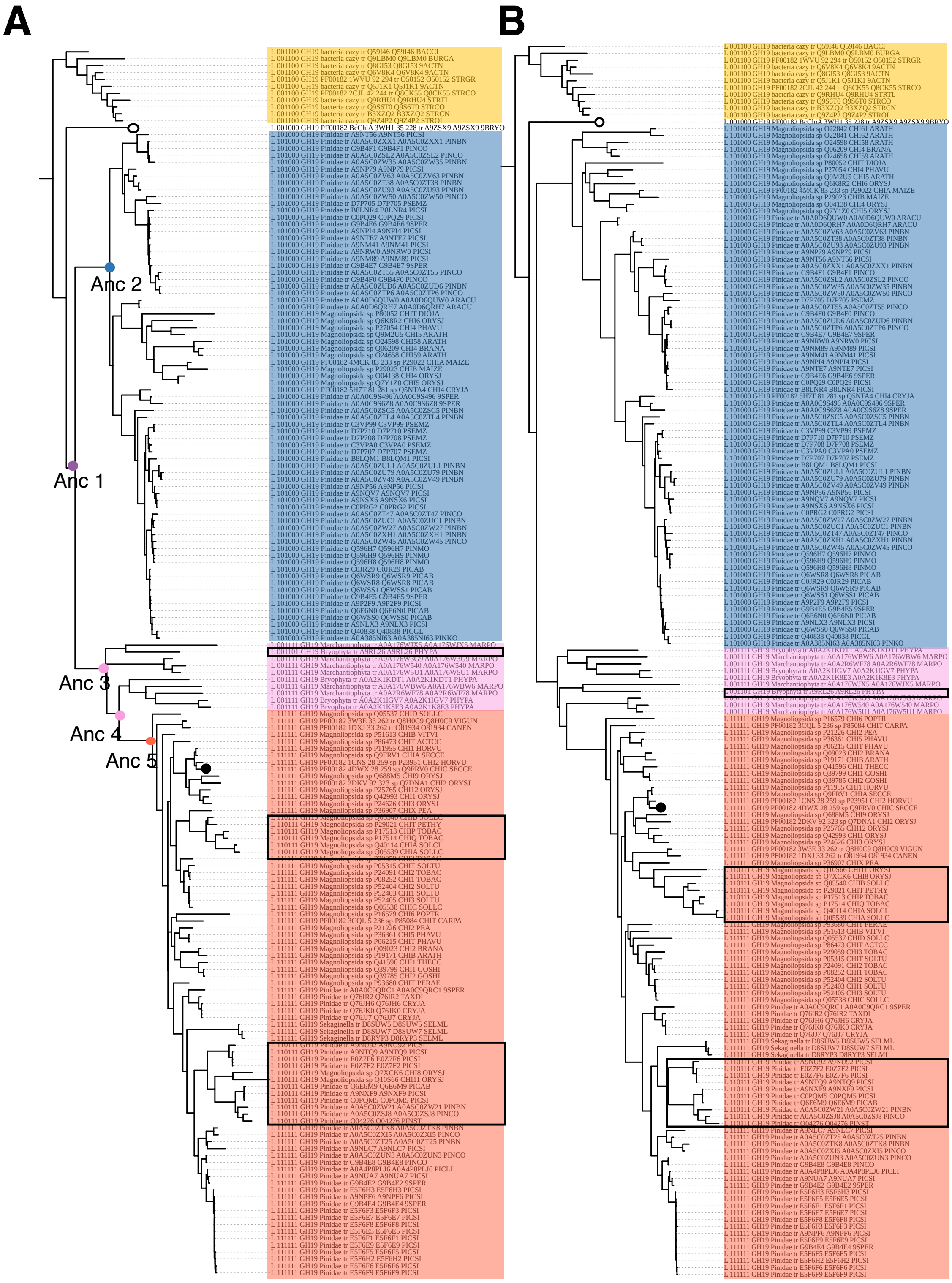
A
B
Anc 1
Anc 2
Anc 3
Anc 4
Anc 5

**Figure S3. Phylogenetic tree of GH19 chitinase constructed with whole sequences of GH19 catalytic domain (A) and sequences without six loop regions (B).**

179 GH19 chitinase sequences containing 168 plants GH19 chitinases and 11 bacterial GH19 chitinases as an outgroup (highlighted in yellow) were used to construct maximum likelihood phylogenetic trees via IQ-TREE using WAG+F+I+G4 model. Sequences with 2, 4, and 6 loops are highlighted in blue, pink, and red, respectively. Open squares indicate the sequences with one loop deletion. The five ancestral nodes that were characterized in this study are labeled and colored according to the number of loop regions (purple, one loop; blue, two loops; pink, four loops; red, six loops).

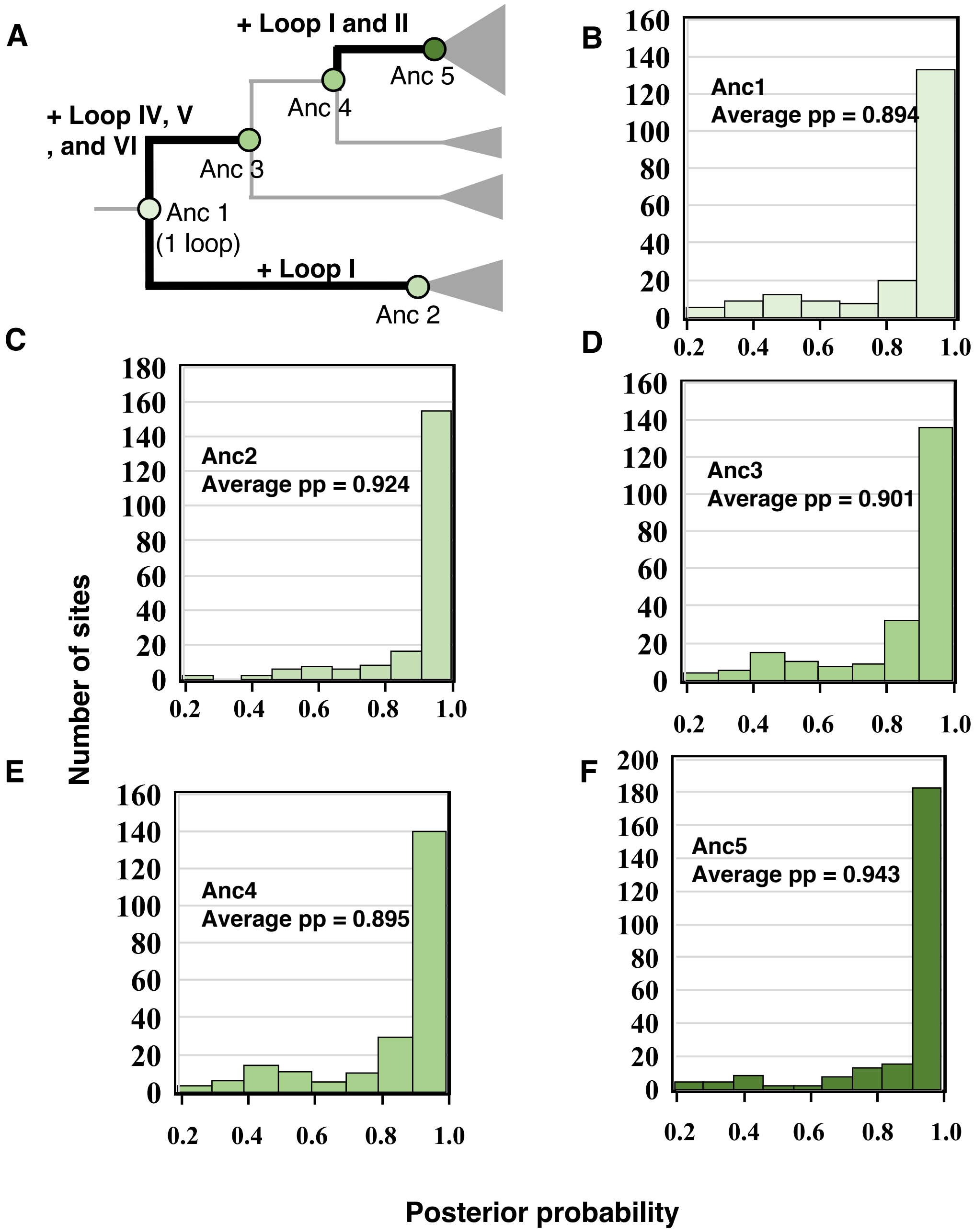

**Figure S4. Site-specific posterior probabilities for ancestral protein sequences.**

A, Schematic representation of a phylogenetic tree of GH19 chitinase.

B-F, Distribution of posterior probabilities (pp) of the maximum a posteriori state at each amino acid site in the inferred ancestral sequences characterized in this study.

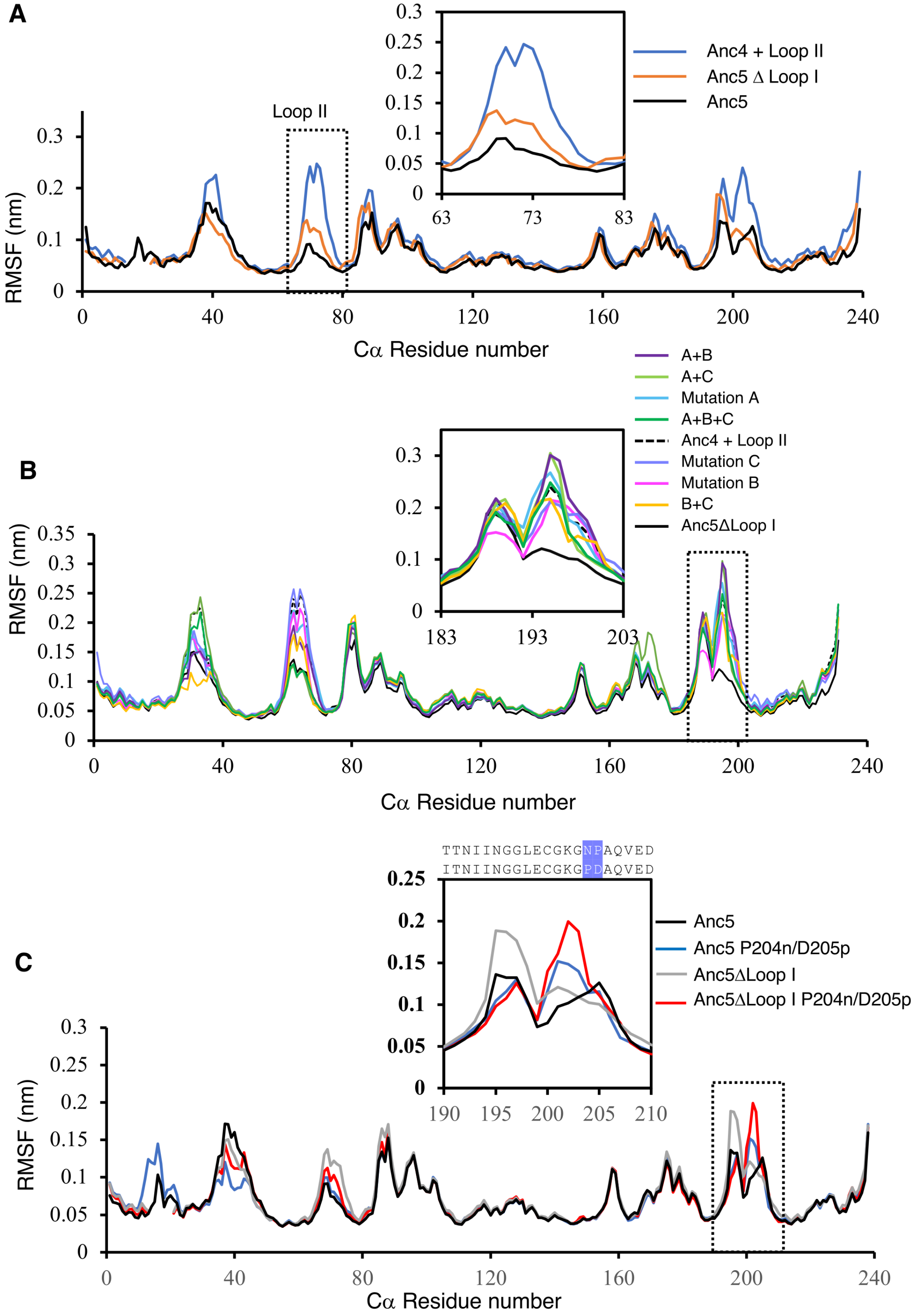
A
Loop II
RMSF (nm)
Cα Residue number
Anc4 + Loop II
Anc5 Δ Loop I
Anc5
B
RMSF (nm)
Cα Residue number
A+B
A+C
Mutation A
A+B+C
Anc4 + Loop II
Mutation C
Mutation B
B+C
Anc5ΔLoop I
C
TTNIINGGLECGKGNPAQVED
ITNIINGGLECGKGPDAQVED
RMSF (nm)
Cα Residue number
Anc5
Anc5 P204n/D205p
Anc5ΔLoop I
Anc5ΔLoop I P204n/D205p

**Figure S5. Mutation effects on loops mobilities.**

A), Plots of the RMSF of Cα of each residue in Anc4 + Loop II (blue), Anc5ΔLoop I (orange), and Anc5 (gray). In the enlargement, it showed reduced mobility of the loop II regions (residue number 66-78) in Anc5ΔLoop I and Anc5 compared to Anc4 + Loop I.

B), Plots of the RMSF of Cα of each residue in Anc5ΔLoop I (black), Anc4 + Loop II (dashed-black), and Anc4 + Loop II mutants. In the enlargement, it showed the movement in a loop region (positions 192 – 201), where the reduced mobility observed in Anc5ΔLoop I compared to Anc4 + Loop II. Introducing substitutions that stabilize loop II and enhance antifungal activity did not stabilize this loop region (positions 192 – 201).

C), Plots of the RMSF of Cα of each residue in Anc5 (black), Anc5ΔLoop I (gray), and their mutants containing Anc4 state (P204n/D205p) in a loop region (positions 199-208, the residue number corresponds to Anc5 state) described in B).

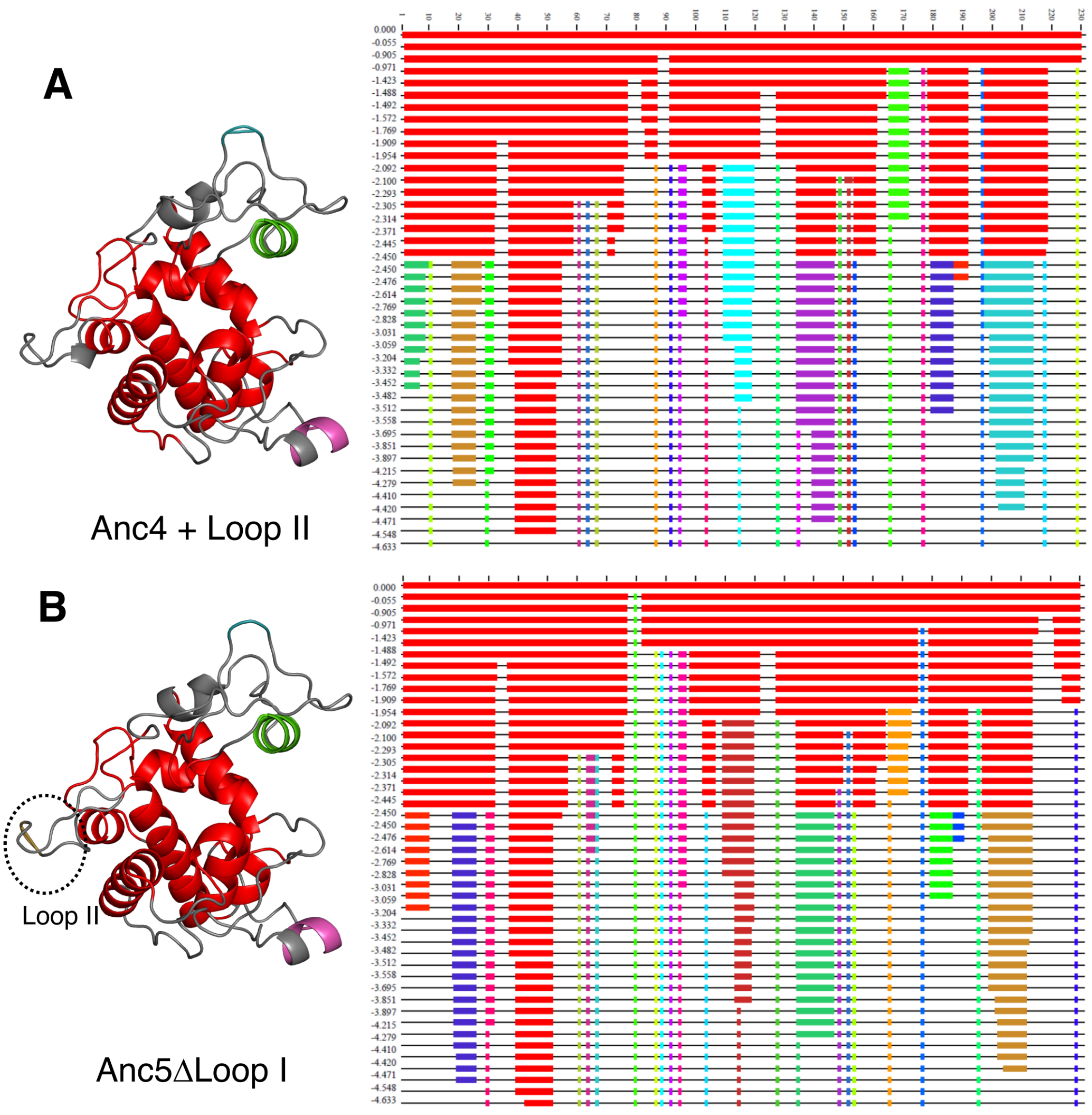

**Figure S6. Computational analysis of long-range communication and rigidity.**

A and B, Left, Rigid cluster decomposition using program FIRST of Anc4 + Loop II and Anc5ΔLoop I at hydrogen bond energy strength cutoff -2.3 kcal/mol are mapped on 3D structures. Distinct rigid clusters are designated by different colored regions, while flexible regions are shown in gray. Red represents the largest dominant rigid cluster. Anc5ΔLoop I has an additional rigid cluster in loop II highlighted with a dashed circle.

Right, Rigidity profile of Anc4 + Loop II and Anc5 ΔLoopI indicated with a hydrogen bond dilution plot with program FIRST. The horizontal axis depicts the residue numbers. The vertical axis represents the current hydrogen bond energy cutoff in kcal/mol. Flexible regions are indicated as black horizontal thin lines and distinct rigid clusters are indicated by various colored blocks with red being the largest rigid cluster. As weak hydrogen bonds are removed, clusters become fragmented, and structure becomes increasingly flexible. Dashed square indicates the rigid cluster within loop II that is observed only in Anc5ΔLoop I.

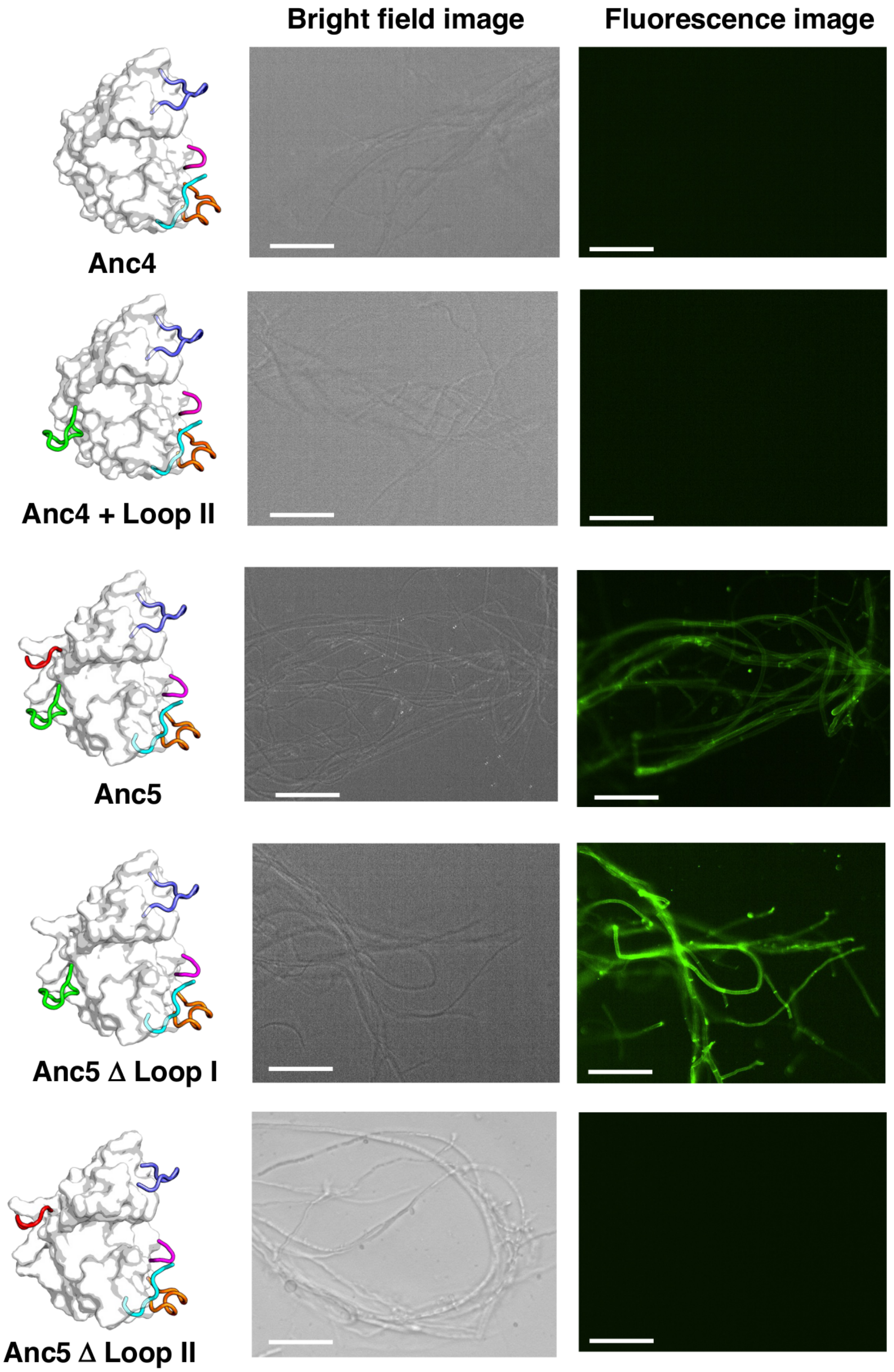
Bright field image
Fluorescence image
Anc4
Anc4 + Loop II
Anc5
Anc5 Δ Loop I
Anc5 Δ Loop II

**Figure S7 Fluorescence microscopy images of fluorophore labelled Anc4 and Anc5 and their mutants on the surface of fungal hyphae revealed that antifungal activity requires the binding activity to the fungal hyphae.**

Each 50 μL of 2 μM Alexa Fluor 488-labelled protein samples are mixed with *T. longibrachiatum* hyphae in 20 mM sodium phosphate buffer, pH 7.4 at 25 °C.

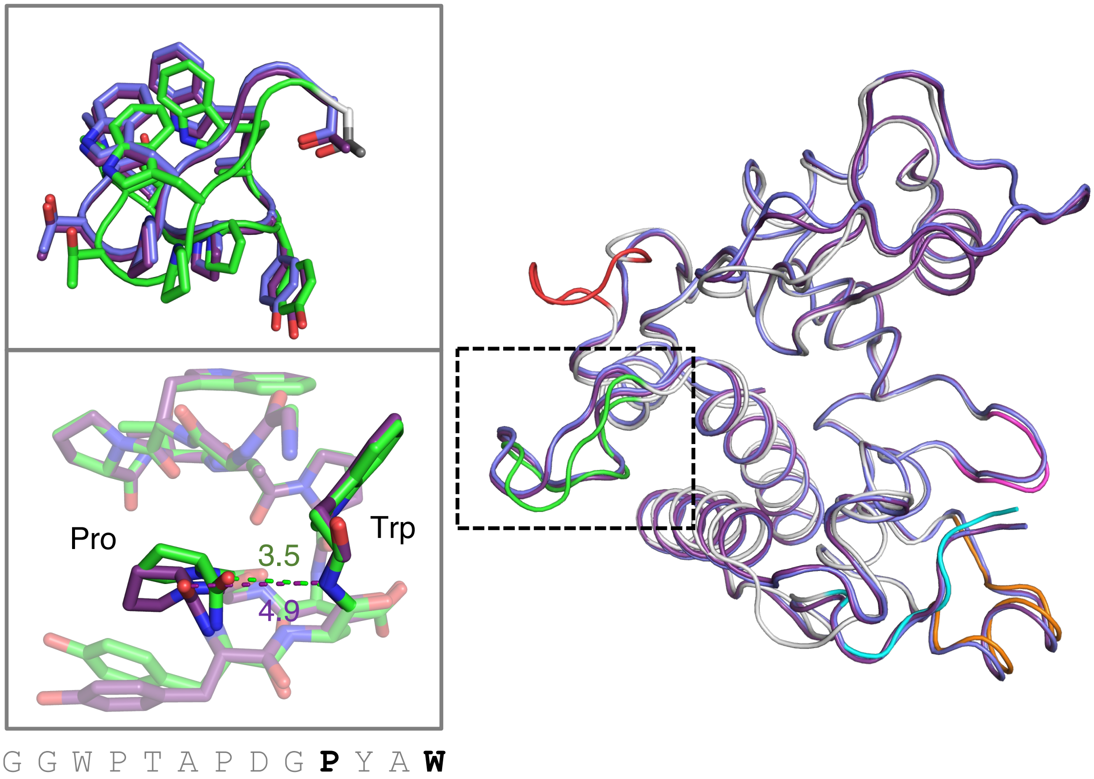

**Figure S8 Structural comparison of Anc5, Anc4+Loop II +A, and Anc4+Loop II +A+B+C.**

Superimpositions of Anc5, Anc4+Loop II +A, and Anc4+Loop II +A+B+C showed that their structures were almost identical with a slight shift in loop II regiosn squared in a dashed line. Cartoon_loop representations of Anc4+Loop II +A and Anc4+Loop II +A+B+C are shown in slate and purple, respectively. Cartoon_loop representations of Anc5 is shown in white with loop regions I-VI highlighted in red, green, slate, magenta, orange, cyan, respectively. In the bottom enlargement, the pairwise structural alignment in loop II regions of Anc5 and Anc4+Loop II +A+B+C revealed that the hydrogen bonding interaction between Pro75 and Trp78 observed in Anc5 (green) was diminished in Anc4+Loop II +A+B+C (purple).
